# Engineering hematopoietic stem and progenitor cells to generate red blood cells as viral traps against HIV-1

**DOI:** 10.1101/2025.04.14.648845

**Authors:** Sofia E. Luna, William N. Feist, Kaya Ben-Efraim, Nelson A. Amorin, Nicole M. Johnston, Hana Y. Ghanim, Freja K. Ekman, Amanda M. Dudek, Matthew H. Porteus

## Abstract

Canonical HIV-1 entry into target cells depends on binding to CD4 as a primary receptor. Because of this, use of the CD4 receptor as a viral trap (a decoy receptor used to prevent infection of target cells) is a promising strategy for the treatment of HIV-1. One challenge in using CD4 viral traps is maintaining enough of the decoy receptor in circulation to remain effective. Here we present a strategy to produce cell-based CD4 viral traps by engineering hematopoietic stem and progenitor cells (HSPCs) to express the CD4 receptor in red blood cell (RBC) progeny. This takes advantage of the ability of the HSPC to repopulate the blood system for a lifetime, while leveraging the fact that RBCs greatly outnumber any cell targeted for infection. Engineered HSPCs efficiently express CD4 on their cell surface after differentiation to the RBC lineage *in vitro*. Fusion of CD4 to glycophorin A (GPA) and introduction of a truncated erythropoietin receptor (tEPOR) leads to increased CD4 expression and enrichment of edited cells (CD4-GPA-tEPOR) to levels capable of neutralizing HIV-1 pseudovirus *in vitro*. In sum, this work presents a potential strategy for the one-time delivery of CD4-RBC viral traps through autologous transplantation of engineered HSPCs.

## Introduction

A key step in viral infection is binding to cell surface receptors that facilitate entry into host cells. Consequently, blocking receptor binding has been a primary focus in the development of new antiviral therapies, which has resulted in the creation of new antibodies, peptides, and small molecules that inhibit receptor and/or co-receptor binding.^1,2^ Another strategy to block viral entry is the use of viral traps, which employ “decoy” receptors that engage with viral particles to prevent binding of target cells. Red blood cells (RBCs) modified to carry decoy receptors represent an intriguing option for the development of viral traps. There are roughly 25 trillion RBCs in adult circulation,^3^ making them the most abundant cell type in the body, vastly outnumbering any other cell type targeted for infection. Moreover, mature RBCs are enucleated and lack the necessary machinery for viral replication, eliminating the potential for viral propagation within RBCs, and are naturally cleared from the body by the reticuloendothelial system.^4,5^ Importantly, naturally occurring RBC viral traps against another virus, Adenovirus Type 5, have been reported, and are hypothesized to have evolved in humans to protect against infection.^6^ Thus, the development of custom RBC viral traps engineered to express decoy receptors builds upon this naturally occurring viral resistance strategy.

One obstacle in utilizing RBCs as viral traps is the difficulty in engineering the enucleated cells to express the receptor of interest. Non-genomic methods of modification have been reported, including adsorption of nanoparticles^7^ and targeting proteins via encapsulation,^8^ antibodies,^9^ and sortase-mediated reactions.^10^ However, these methods are limited by the short survival of the modified cells in circulation (maximum reported ∼28 days) due both to dissociation of the modifications and natural clearance of the cells.^10^ An alternative strategy is genetic engineering of erythroid progenitor cells and subsequent differentiation into enucleated RBCs.^11^ This approach offers a permanent alteration of engineered RBCs, yet transfused RBCs only last around 50-60 days^12^ and therefore require repeated administration to combat chronic infections. Here we present a strategy to overcome these obstacles by genome-engineering hematopoietic stem and progenitor cells (HSPCs) for erythroid-specific expression of CD4 to generate CD4-expressing RBC viral traps against HIV-1.

HIV-1 infects cells by binding to the CD4 cell-surface receptor, leading to a conformational change that allows for binding to one of two co-receptors, CCR5 or CXCR4, followed by fusion with the host cell membrane.^13^ Although co-receptor switching and drug resistance mutations are common, all strains of HIV-1 rely on binding to the CD4 receptor for infection. Consequently, viral mutations to escape binding will also lead to an accompanying decrease in viral fitness.^14,15^ Thus, prior work has sought to capitalize on this reliance by repurposing exogenous CD4 as a decoy to prevent binding to target cells (primarily CD4^+^ T cells and macrophages). Early works employed delivery of a soluble form of CD4 (sCD4), which successfully neutralized laboratory strains of HIV-1, but showed a limited efficacy in clinical trial.^16^ An improvement on this strategy involved combining CD4 with an immunoglobulin Fc region (CD4-Ig)^17^ and a CCR5-mimetic sulfopeptide (eCD4-Ig),^18^ of which the latter protected rhesus macaques against simian/human immunodeficiency virus (SHIV) challenges.^18,19^ A recent strategy employed CD4-expressing virus like particles (VLPs),^15^ and, later, RBCs,^11^ which allowed for formation of CD4 receptor clusters on the VLP or RBC surface. These clusters mimic natural high avidity binding seen on cell surfaces, thereby mitigating the potential for viral escape.^11,15^ Using a cell-based carrier to deliver CD4 has the potential to dramatically improve the half-life of delivered drugs,^5^ yet these cells will eventually require re-administration. Despite great advancement in the efficacy of CD4 mimetic compounds, they all require repeated dosing to remain effective due to natural clearance from circulation. In contrast, transplantation of engineered HSPCs presents a one-time strategy for lifelong production of RBC viral traps.

In this study, we engineer HSPCs so that upon reconstitution of the hematopoietic system they will generate CD4-expressing RBCs that serve as viral traps against HIV-1. Key to our strategy is precisely editing HSPCs for erythroid-specific expression, so that only derived RBCs and no other blood cell lineages express CD4. This lineage specific expression harnesses the critical properties of RBCs to serve as highly abundant viral traps, while leaving other lineages unperturbed. We demonstrate efficient engineering of HSPCs that successfully differentiate into CD4-expressing RBCs *in vitro*. We also introduce an RBC-specific selective advantage in the edited cells through introduction of a truncated erythropoietin receptor (tEPOR)^20^ and show that derived RBCs significantly reduce viral infection *in vitro*. Taken together, we introduce a strategy for inhibition of chronic viral infection through autologous transplantation of HSPCs engineered for RBC-specific expression of viral receptors.

## Results

### Precision integration of CD4 expression cassettes in human HSPCs

Recent studies have shown that CD4 decoys, such as CD4-expressing RBCs and eCD4-Ig, can be used to efficiently neutralize HIV infection.^11,15,18,19,21^ Based on this concept, we hypothesized that knock-in of a CD4 expression cassette into the *HBA1* locus of HSPCs would allow for high levels of lineage-specific production of CD4-expressing RBC progeny to act as viral traps, while minimizing the potential of CD4 expression in other blood cell types. When we analyze the expression of *HBA1* in the Human Protein Atlas database, we find it is only expressed in RBCs and has a mean expression level of >50,000 normalized transcripts per million (nTPM), among the highest expressed genes in any cell type^22,23^. Targeting *HBA1* functions as a safe harbor integration due to the natural redundancy of *HBA1* with *HBA2* and the fact that individuals with bi-allelic inactivation of *HBA1* have the benign condition of α-thalassemia trait. While the random integration of AAV is rare because it contains no integrase, the design of the construct without a promoter also limits any potential for toxicity from an off-target integration event (such as insertional oncogenesis as seen in gene therapy trials for SCID-X1, chronic granulomatous disease, and X-linked adrenoleukodystrophy). We used ribonucleoprotein (RNP) based CRISPR/Cas9 editing along with recombinant AAV6 delivered donor templates to target CD4 expression cassettes into the *HBA1* locus by whole gene replacement.^24^ We employed a previously described single-guide RNA (sgRNA) that specifically cuts in the 3’ untranslated region (UTR) of *HBA1* while leaving *HBA2* intact, and designed donor templates with homology arms flanking the coding sequence of *HBA1* (Fig. 1A).^24^ Based on prior work by Hoffman et al. (2021), who used semi-random integrating lentiviral integrations,^11^ we tested two forms of CD4 donor templates: the entire CD4 coding sequence (codon optimized and hereafter termed “CD4”) and a CD4-GPA fusion protein, which physically links the codon-optimized extracellular D1D2 region of human CD4 to the N-terminus of glycophorin A (GPA) (hereafter termed “CD4-GPA”). This earlier study found that codon-optimization of CD4 leads to increased expression of the receptor, and that use of CD4-GPA increased CD4 expression on erythrocytes since GPA is naturally highly expressed on the surface of RBCs.^11^ We also designed two additional vectors containing a truncated erythropoietin receptor (tEPOR) following a 2A self-cleavage peptide on the 3’ end of either CD4 or CD4-GPA (Fig. 1A). Use of tEPOR has been shown to increase erythroid proliferation in an erythropoietin-dependent manner and enrich for tEPOR-edited cells over the course of erythroid differentiation.^25,26^

**Figure 1:**
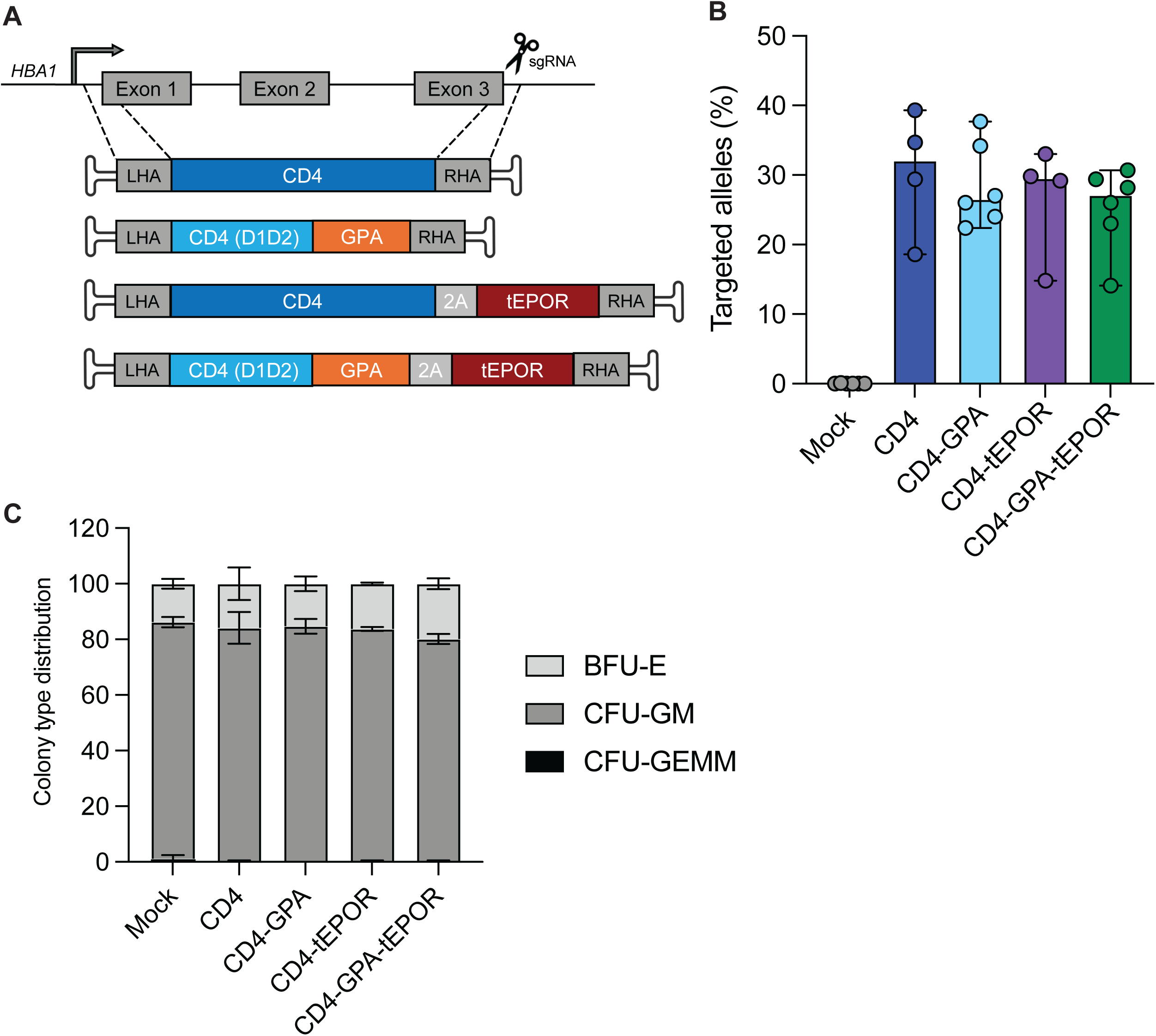
Efficient engineering of CD4 receptor cassettes at the *HBA1* locus. **A.** Schematic of genomic editing strategy at the *HBA1* locus for targeted integration of CD4 expression cassettes delivered by AAV6. **B.** Percentage of alleles with targeted integration for each CD4-expression cassette. Each AAV6 was used at 2500 vector genomes/cell. n=6 independent HSPC donors for Mock, CD4-GPA, CD4-GPA-tEPOR, n=4 for CD4 and CD4-tEPOR. Bars represent median +/− 95% confidence interval. **C.** CFU assay showing mock edited cells versus cells edited with each CD4 expression cassette. Bars represent relative frequency of each progenitor colony type: CFU-GEMM (multi-potential granulocyte, erythroid, macrophage, megakaryocyte progenitor cells), CFU-GM (colony forming unit-granulocytes and monocytes), BFU-E (erythroid burst forming units) (n=3 independent HSPC donors). Bars represent mean +/−SD.

To test targeted integration frequency of the four CD4 donor cassettes, we delivered high fidelity Cas9 protein complexed with the *HBA1* sgRNA into HSPCs via electroporation, followed by immediate transduction with individual AAV6 donors. Three days post-editing, we analyzed the targeted integration frequency at *HBA1* by droplet digital PCR (ddPCR). After optimization of AAV6 multiplicity of infection (MOI) (Supplementary Fig. 1A), we achieved an average allelic integration frequency of approximately 30% using 2500 vector genomes/cell (vg/cell) (Fig. 1B). To assess the impact of the editing on the differentiation capacity of HSPCs, we performed a colony forming unit (CFU) assay and found that while there was a decrease in total colony formation consistent with prior reports of RNP/AAV6-mediated editing (Supplementary Fig. 1B),^27–31^ the lineage distribution of colonies was not affected, indicating that targeted integration of the CD4 expression cassettes does not impede myelo-erythroid differentiation potential of HSPCs *in vitro* (Fig. 1C). Additionally, when we repeated the assay with higher purity AAV6 vector isolated by cesium chloride (CsCl) gradient ultracentrifugation, there was a marked increase in relative total colony formation in the genome-edited conditions, with the distribution of lineage formation once again remaining the same (Supplementary Fig. 1C). When using AAV6 for HDR in HSPCs, the quality of the AAV is of critical importance and these results are consistent with previous reports showing that high quality AAV6 preparations allow for better HSPC function.^32^

### Edited HSPCs differentiate into CD4-expressing red blood cells *in vitro*

Next, the HSPCs targeted with each CD4 expression cassette were differentiated into RBCs using a previously described 14-day *in vitro* erythroid differentiation protocol (Fig. 2A).^33^ On day 14, we assessed erythroid differentiation by flow cytometry using established RBC markers (CD34^-^CD45^-^ CD71^+^). We observed no appreciable differences in phenotypic RBC differentiation markers between mock-edited and CD4-edited cells. On average >85% of cells in all conditions expressed CD71 on day 14, indicating that addition of CD4-expression cassettes does not impede RBC differentiation (Fig. 2B). Of note, GPA, a commonly used RBC cell surface marker, was omitted during quantification of RBC differentiation due to antibody cross-reactivity with the GPA expressed in some of the editing conditions. However, we found no differences in GPA expression in the CD4 and CD4-tEPOR edited conditions compared to mock edited cells (Supplementary Fig. 2A). Additionally, when we looked for more mature markers of RBC formation on day 14 and 18 of differentiation, we found that mock, CD4-GPA and CD4-GPA-tEPOR edited cells were all able to gain Band3 expression and lose α4-integrin expression as expected for normal RBC differentiation (Supplementary Fig. 2B).

**Figure 2:**
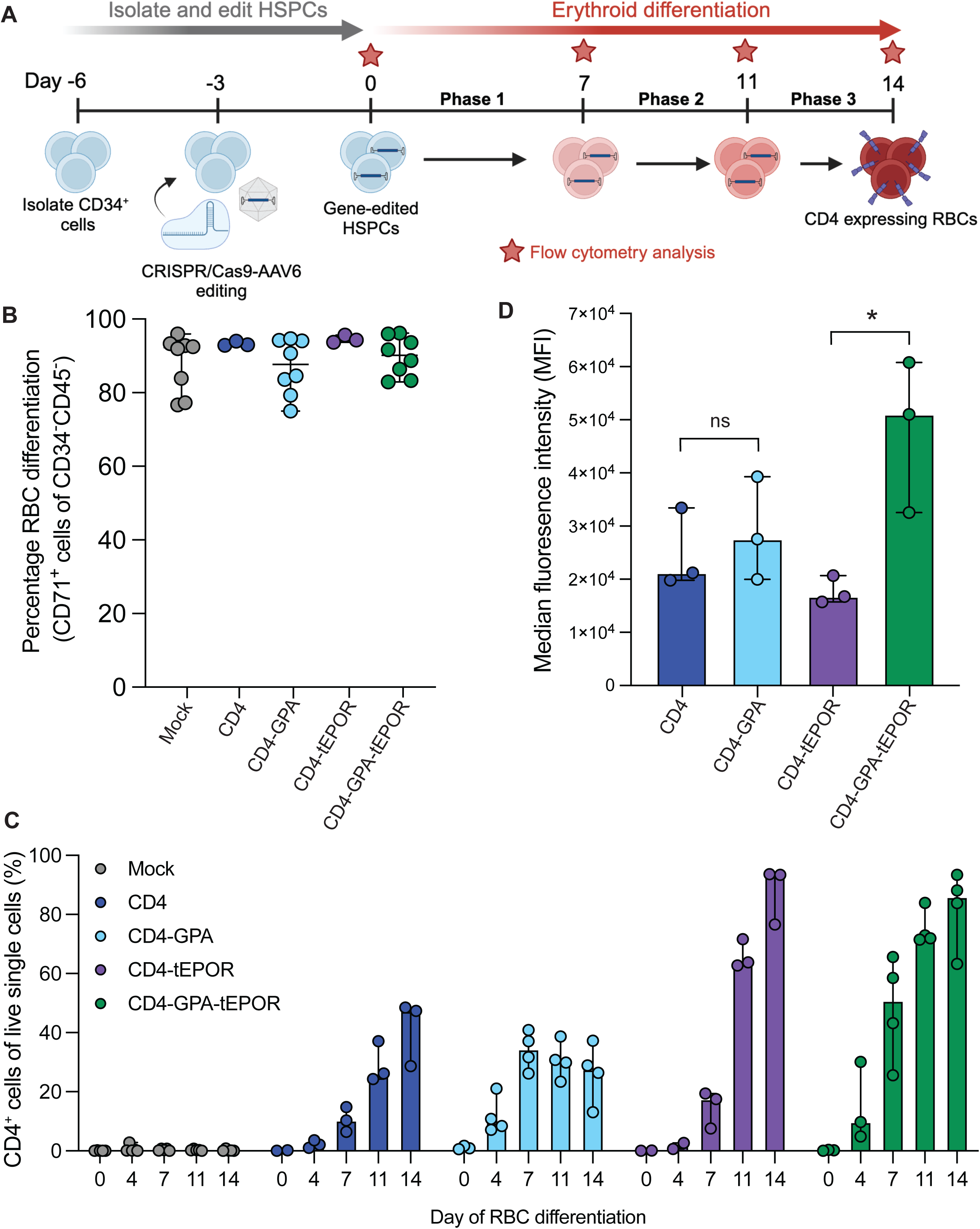
Genome-edited HSPCs show robust expression of CD4 following erythroid differentiation. **A.** Diagram showing workflow of genome-editing and subsequent RBC differentiation of HSPCs. **B.** Percentage of CD71^+^ cells of CD34^-^CD45^-^ cells on day 14 of RBC differentiation. n=8 for Mock, CD4-GPA, CD4-GPA-tEPOR, n=3 for CD4 and CD4-tEPOR. **C.** Percentage of CD4^+^ cells of live single cells on each day of RBC differentiation. n=6 for Mock, n=4 CD4-GPA, CD4-GPA-tEPOR, n=3 on day 4-14 for CD4 and CD4-tEPOR on day 4-14 and n=2 on day 0. **D.** Median fluorescence intensity (MFI) of CD4 signal within CD4^+^ cells on day 14 of RBC differentiation. n=3 for all conditions. ns, p=0.5919 for CD4 vs. CD4-GPA, *p=0.0224 for CD4-tEPOR vs. CD4-GPA-tEPOR by unpaired two-tailed *t*-test. All bars represent median +/−95% confidence interval and all replicates represent independent HSPC donors.

After confirming there was no defect in differentiation, we quantified relative CD4 receptor expression on the surface of cells throughout differentiation in each editing condition. As expected, because CD4 expression is driven by the RBC-specific endogenous *HBA1* promoter, there were no CD4^+^ cells in any of the conditions at the beginning of differentiation (termed day 0). By the end of RBC differentiation (day 14), an average of 85% of cells expressed CD4 in the CD4-tEPOR and CD4-GPA-tEPOR conditions, while an average of 33% of cells displayed CD4 in the constructs lacking tEPOR (Fig. 2C). As we hypothesized, by expressing the protein from the robust *HBA1* promoter in the RBC lineage, we found that CD4 was highly expressed on the cell surface at day 14 (Supplementary Fig. 3A). Additionally, we found that CD4 expression was indeed RBC-specific and only detectable in cells expressing the erythroid marker CD71 (Supplementary Fig. 3B). These data highlight that our knock-in expression constructs show high levels of CD4 expression, are highly lineage specific, and the total number of CD4^+^ cells is augmented by the inclusion of tEPOR.

Allelic integration frequencies of the CD4 expression cassettes at the *HBA1* locus were monitored by ddPCR throughout differentiation and confirmed that targeted integration frequencies correlate with CD4 cell surface expression (Supplementary Fig. 4A). As previously shown in other tEPOR-based expression systems^20^, CD4-tEPOR and CD4-GPA-tEPOR edited cells enriched over the course of differentiation, achieving an average targeted integration frequency of almost 70% on day 14. This represented an average fold increase of 2.8x for CD4-tEPOR and 2.9x CD4-GPA-tEPOR from day 0 to day 14 of differentiation. The knock-in rate of non-tEPOR constructs remained at around 30% throughout differentiation (Supplementary Fig. 4A). Because we observed robust enrichment over time from an already high knock-in frequency, we then investigated how strong the enrichment phenotype could be under lower efficiency editing conditions. We edited HSPCs with half the AAV6 MOI (1250 vg/cell) using the CD4-tEPOR and CD4-GPA-tEPOR cassettes to evaluate the extent to which tEPOR-edited cells could enrich over the course of RBC differentiation. While initial editing frequencies were indeed lower at day 0 (average 18.5% for CD4-tEPOR and 17.2% CD4-GPA-tEPOR), edited cells enriched further throughout differentiation (average fold increase of 3.8x for CD4-tEPOR and 3.5x CD4-GPA-tEPOR from day 0 to day 14), reaching similar integration frequencies to those observed in the cells edited with the higher AAV6 dosage by the end of RBC differentiation (Supplementary Fig. 4B).

Lastly, we examined the abundance of CD4 receptors on the cell surface by measuring the median fluorescence intensity (MFI) of CD4 signal by flow cytometry. We found both CD4-GPA and CD4-GPA-tEPOR constructs exhibited higher levels of CD4 receptor expression when compared to the CD4 and CD4-tEPOR constructs, respectively, with CD4-GPA-tEPOR yielding the highest MFI (Fig. 2D). We then compared the relative abundance of CD4 receptors on the CD4-GPA and CD4-GPA-tEPOR RBCs compared to stimulated and unstimulated CD4^+^ T cells and found that the CD4-expressing RBCs had 2 to 10-fold higher expression of CD4 receptors on the cell surface (Supplementary Fig. 4C). Using a 2-fold higher expression (the lower end of what we find) and the relative abundance of RBCs compared to CD4^+^ T-cells (roughly 50 times more), if only 1% of RBCs were derived from engineered HSPCs, there would still be 100 times more expression of CD4 on RBCs than on T cells. This would create a hostile environment for HIV that we would expect to lead to the virus being slowly extinguished from the body. These results demonstrate the strong potential of CD4-engineered RBCs to act as viral traps for HIV-1 due to both greater expression of the CD4 on the cell surface and their naturally greater abundance in circulation compared to T cells.

### CD4-GPA expressing myelo-erythroid cell line neutralizes HIV-1 pseudovirus

To investigate whether the cells edited with the full-length CD4 cassette or CD4-GPA fusion cassette have superior neutralization capacity, we tested the potential of cells expressing each construct to neutralize HIV-1 pseudovirus using a CD4 or CD4-GPA expressing K562 (myelo-erythroid) stable cell line. The cell lines were created by targeting cassettes carrying each receptor (CD4 or CD4-GPA) driven by a strong ubiquitous promoter, Ef1α, and co-expressed with green fluorescent protein (GFP) to the *CCR5* safe-harbor locus (Fig. 3A). RNP targeting *CCR5*^34^ was nucleofected into K562 cells along with a CD4-2A-GFP or CD4-GPA-2A-GFP plasmid. We then used fluorescence activated cell sorting (FACS) to isolate a pure population of GFP^+^ cells. Post-sorting, we confirmed that both CD4 and CD4-GPA K562 cell lines contained >90% CD4^+^ cells (Fig. 3B). We found that the CD4-GPA expressing cells exhibited much higher levels of expression of the CD4 receptor on the cell surface, with an MFI almost 5 times higher than its CD4 counterpart (Fig. 3C-D). To test the ability of the cells to neutralize HIV-1, we employed a modified version of a TZM-bl neutralization assay adapted from a previous report using NL4-3ΔEnv-GFP pseudotyped with a clade B envelope, TRO11.^11,35^ At all cell concentrations tested, only the CD4-GPA expressing cells neutralized the HIV pseudovirus (Fig. 3E). While neutralization was only observed at the highest concentration of 6×10^7^ cells/mL, the concentration of circulating RBCs in the blood is almost 100-times greater than this (∼5×10^9^ cells/mL),^36^ suggesting edited RBCs may serve as effective viral traps *in vivo*. Overall, these results indicated that only the CD4-GPA cells drove sufficient CD4 receptor expression on the cell surface to facilitate viral entrapment, therefore we chose to proceed in future experiments with only the CD4-GPA constructs.

**Figure 3:**
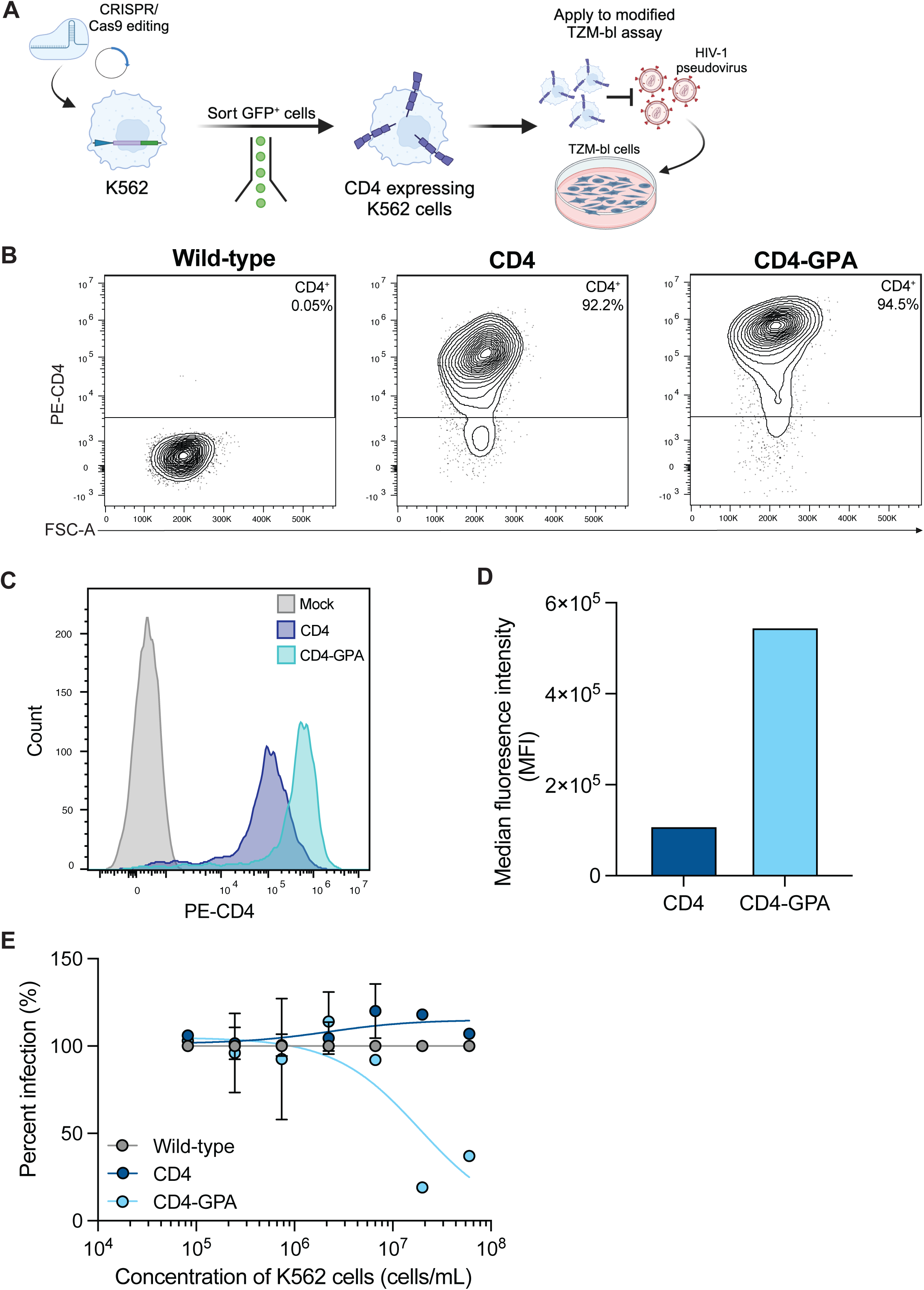
Stable cell line expressing CD4 neutralizes HIV-1 pseudovirus. **A.** Schematic of workflow for K562 stable cell line generation and testing. **B.** Flow cytometry plots of CD4 expression on wild-type K562s and K562s engineered with CD4 expression cassettes driven by the Ef1α promoter after sorting and expansion of GFP^+^ cells. **C.** Histogram showing relative CD4 expression of wild-type, CD4, and CD4-GPA K562s. **D.** Median fluorescence intensity (MFI) of CD4 signal within CD4^+^ cells for CD4 versus CD4-GPA K562s. n=1. **E.** Inhibition of infection with TRO11 HIV-1 pseudovirus using wild-type, CD4, or CD4-GPA engineered K562s (n=1, data points and error bars represent mean +/−standard deviation of technical duplicate infections, some duplicates were excluded due to mechanical error).

### HSPC-derived CD4^+^ red blood cells neutralize HIV-1 pseudovirus

To determine whether HSPC-derived CD4^+^ RBCs were able to neutralize HIV-1 we employed the same editing and erythroid differentiation protocol as above and then applied the derived RBCs to the modified TZM-bl neutralization assay (Fig. 4A). Briefly, CD34^+^ HSPCs were targeted with either CD4-GPA or CD4-GPA-tEPOR followed by RBC differentiation. On day 14, the ability of cells to neutralize HIV-1 was evaluated through a modified TZM-bl assay using TRO11 pseudovirus. In three assays performed with independent biologic HSPC donors, the CD4-GPA-tEPOR RBCs reduced viral infection at the highest cell concentration tested of 6×10^7^ cells/mL (Fig. 4B). In contrast, CD4-GPA RBCs only neutralized viral infection in one of the three donors (Donor 1) at a concentration of 6×10^7^ cells/mL (Fig. 4B). None of the lower cell concentrations were able to inhibit infection in cells edited with either construct. On average, CD4-GPA-tEPOR RBCs were able to inhibit 52% of HIV pseudovirus infection at the highest cell concentration tested when normalized to mock edited cells, while CD4-GPA RBCs inhibited less than 10% of infection (Fig. 4C). This difference in the neutralization capacity is likely due to the much greater number of CD4^+^ RBCs generated in the CD4-GPA-tEPOR condition due to enrichment in edited cells driven by the tEPOR (average 27.2% of RBCs in CD4-GPA versus 80.3% in CD4-GPA-tEPOR) (Supplementary Fig. 5A-B). Because clade B envelopes such as TRO11 tend to be easily neutralized, we also tested edited RBC neutralization capacity against a clade CRF01 envelope, CNE55,^35^ and found similar neutralization capacity as with TRO11 (Fig. 4D). This highlights the fact that our strategy is designed to be agnostic to HIV-1 clade or subtype, due to the reliance of all HIV-1 on CD4 binding for infection. Importantly, the highest concentration of CD4-RBCs tested in these experiments (6×10^7^ cells/mL) is equivalent to just ∼1.2% of the RBC concentration in human circulation (∼5×10^9^ cells/mL).^36^ Therefore, these results display the potential of CD4-GPA expressing RBCs to efficiently neutralize virus *in vivo,* even if only a fraction of edited cells engraft and repopulate the blood system.

**Figure 4:**
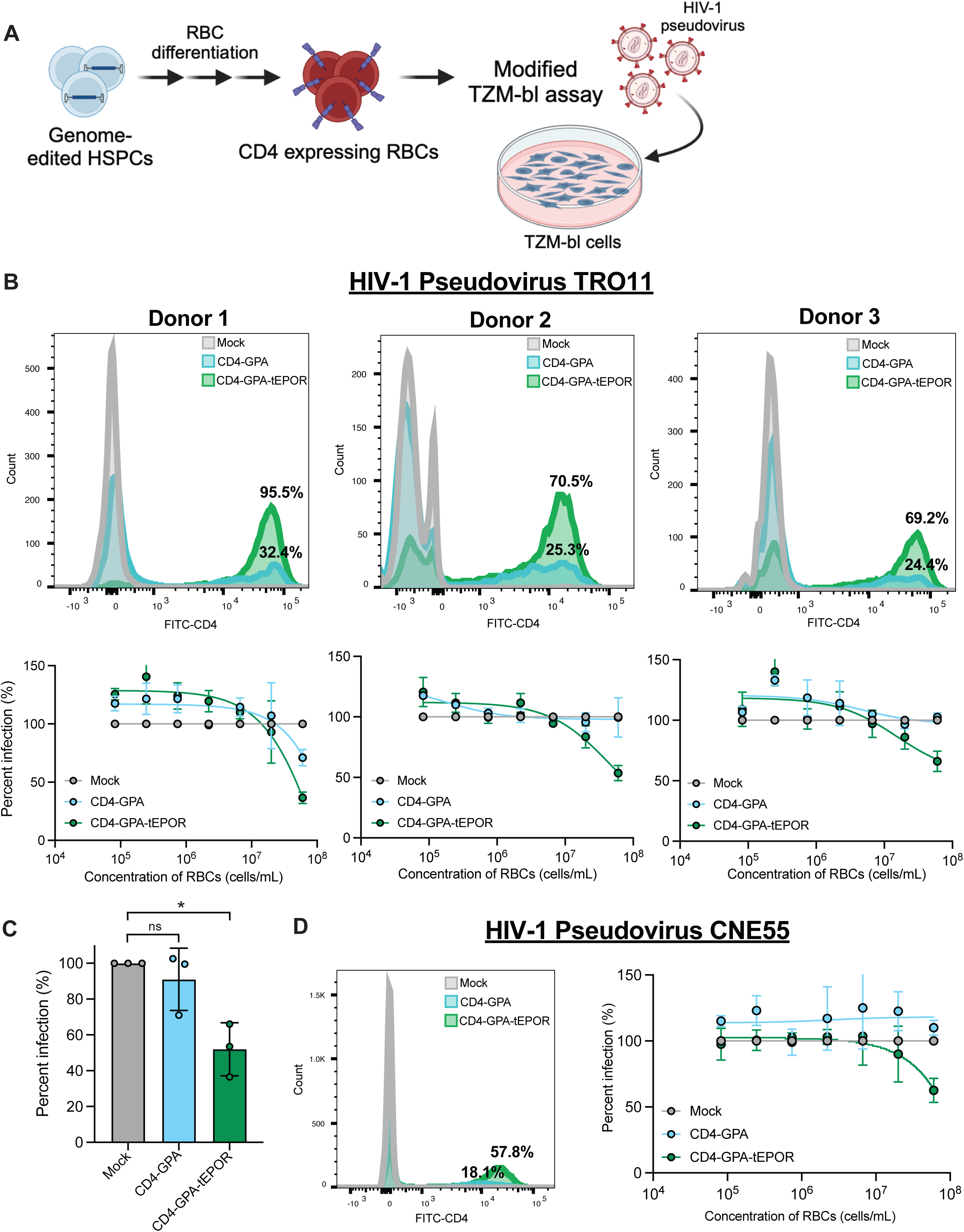
CD4 expressing RBCs neutralize HIV-1 pseudovirus. **A.** Schematic of genome-editing of HSPCs followed by RBC differentiation and application to the modified TZM-bl assay. **B.** Inhibition of infection with TRO11 HIV-1 pseudovirus by differentiated mock or CD4-expressing RBCs of each condition as indicated. Histogram panels (top) show relative CD4 expression of live single cells from each condition and biological donor. Bottom panels show percent infection at each concentration of gene-edited RBCs normalized to infection at each concentration of mock edited RBCs. n=3 biological donors shown in separate panels. Data points and error bars represent mean +/− standard deviation of technical duplicate infections. **C.** Percent infection of TRO11 HIV-1 pseudovirus at the highest cell concentration tested (6×10^7^ cells/mL) when normalized to mock edited RBCs. n=3 biological donors also shown in panel B. Bars represent mean +/− standard deviation of three independent biological donors. ns, p = 0.4206 for Mock vs. CD4-GPA, *p=0.0049 for Mock vs. CD4-GPA-tEPOR by unpaired two-tailed *t*-test. **D.** Inhibition of infection with CNE55 HIV-1 pseudovirus by differentiated mock or CD4-expressing RBCs of each condition as indicated. Histogram panel (right) shows relative CD4 expression of live single cells in each condition. Right panels show percent infection at each concentration of cells from genome-engineered RBCs normalized to infection at each concentration of mock edited RBCs. Values normalized to infection at each concentration of mock edited RBCs. n=1 biological donor. Data points and error bars represent mean +/− standard deviation of technical duplicate infections (three mock duplicates were excluded due to mechanical error).

## Discussion

We present a novel precision genome-editing strategy of HSPCs to produce CD4-expressing RBC viral traps as a treatment for HIV-1 through autologous HSPC transplantation. There is precedent for using HSPC transplantation to treat HIV-1 as the only reported long-term cure of HIV-1 infection is via allogeneic transplantation of *CCR5*-null HSPCs.^37–39^ While this is a promising strategy, allogeneic transplantation is not a viable treatment option for most patients due to both the high risks associated with allogeneic transplantation, most notably graft versus host disease (GvHD), and the scarcity of matched donors carrying the *CCR5*-null mutation. Additionally, loss of CCR5 alone is ineffective for patients who carry CXCR4- or dual-tropic HIV-1.^40,41^ The strategy presented here overcomes each of these barriers. First, engineering and transplanting a patient’s own cells via autologous transplantation removes the need for a matched donor and decreases the risks associated with allogeneic transplantation. Second, engineering HSPCs to express the CD4 receptor on RBC progeny of transplanted cells represents a global strategy which is agnostic to co-receptor tropism or HIV-1 clade, as all HIV-1 strains must use the CD4 receptor to infect cells despite the high global diversity of circulating HIV-1 envelope glycoproteins. Altogether, the work presented here marks a first step toward the development a one-time strategy for functional cure of HIV-1 that builds on the past success of HSPC transplantation.

In this study, we demonstrate that knock-in of CD4 cassettes in HSPCs for RBC-specific expression can produce effective viral traps against HIV-1. First, we show efficient knock-in of CD4 expression cassettes at the *HBA1* locus in HSPCs using an RNP/AAV6-mediated whole-gene replacement strategy to tie expression to the endogenous *HBA1* promoter (Fig. 1). By using the *HBA1* promoter, we restrict the expression of CD4 to only the RBC lineage, minimizing perturbation of other cell lineages and taking advantage of the natural properties of RBCs to serve as viral traps. We then show that when engineered HSPCs differentiate into RBCs *in vitro,* they express CD4 on the cell surface (Fig. 2). Aligning with prior work,^11^ we also demonstrate that combining the CD4 with GPA, an abundant RBC cell surface protein, allows for greater expression of CD4 on the cell surface (Fig. 2 & 3). Moreover, pairing CD4 expression with that of a truncated erythropoietin receptor (tEPOR) leads to increased production of CD4-RBCs, allowing for approximately 80% of resultant cells to express CD4 by the end of differentiation (Fig. 2). Lastly, we show that CD4-GPA-tEPOR edited RBCs can neutralize HIV-1 pseudoviral infection *in vitro*, while CD4-GPA edited RBCs only showed minimal neutralization in one donor, likely due to the disparity in the number of CD4-expressing cells between conditions (Fig. 4). Because there is roughly a 100-times greater concentration of RBCs in human circulation than we were able to recapitulate *in vitro*, we believe this strategy has the potential to neutralize HIV-1 *in vivo*. With this in mind, we can perform a rough calculation of the level of engineered HSPC engraftment that would be necessary to achieve neutralizing levels of RBC viral traps in circulation. Considering that the incorporation of the tEPOR provides at least a three-fold advantage to knock-in cells during RBC development, it is possible that engraftment as low as 3-4% could result in a 10% circulating concentration of engineered RBCs (a concentration approximately 10 times higher than the levels tested *in vitro*). Such a low level of engineered cell chimerism might be achievable with non-myeloablative, non-chemotherapy based conditioning regimens prior to autologous cell transplantation, making this therapy both safer and more accessible for patients.

While this work provides a promising proof-of-concept for HSPC-derived RBC viral traps, there are several areas that remain to be explored before this strategy moves toward the clinic. Most notably, because it is well-known that immunodeficient mice transplanted with human HSPCs do not reconstitute with many human RBCs in circulation due to phagocytosis by mouse macrophages,^42,43^ we were not able to evaluate the *in vivo* production of CD4-expressing RBCs in a mouse model. However, previous studies have demonstrated that HSPCs edited to induce HIV resistance are able to facilitate multi-lineage engraftment of human cells *in vivo*.^44,45^ Future preclinical work will need to define an *in vivo* model to study how long engineered RBCs persist in circulation and how they are cleared. In particular, it will be important to identify whether engineered RBCs are cleared more quickly through the reticuloendothelial system, or if there may be other undesirable clearance pathways of the RBC viral traps, such as through engulfment by macrophages. Future works may look to HSPC transplantation in the non-human primate (NHP) model for *in vivo* characterization of engraftment of modified cells and subsequent engineered RBC persistence and clearance. In addition, this would allow for investigation of the efficacy of our transplant therapy, as NHPs can be challenged with SHIV.

The use of CD4 viral traps exploits a major vulnerability of HIV-1 because all strains of HIV-1 bind CD4 to enter target cells, thus mutations to escape binding lead to a decrease in viral fitness^14,15,46,47^. Moreover, because CD4 is naturally found in clusters on the surface of T cells and macrophages^48,49^, HIV-1 Env binds to multiple CD4 receptors (leading to avidity^15,50^) in order to infect cells^51,52^. By more accurately mimicking the presentation of CD4 on the HIV-1 target cell through presentation on the RBC cell surface, we decrease the likelihood of viral escape seen when using a monomeric forms of CD4^15,53^. Despite these advantages, it is well-known that it is important to use multiple means of resistance to combat HIV-1 and the strategy presented here could be readily combined with other CRISPR-Cas9 based strategies being developed by our group and others to engineer HIV-1 resistance, such as *CCR5* KO alone^54,55^ or in combination with knock-in of either HIV-1 inhibiting proteins^34^ or neutralizing antibodies.^45^ We envision a future where multiple genome-editing strategies could be combined in order to produce HSPCs with potent multilayered resistance against HIV-1, which is essential to combating a virus well-known to escape single forms of resistance. In particular, the strategy presented here is especially poised to combine well with KO or knock-in strategies to *CCR5,* as it functions by knocking into the RBC-specific safe harbor locus *HBA1*, therefore not competing for the same genomic locus. Our group and others have demonstrated feasibility of multiplexed editing at multiple genomic loci in primary cells,^20,56,57^ and as this approach improves in the future, engineering resistance at both the *CCR5* and *HBA1* loci has the potential to deliver cells with broader resistance against HIV-1.

Overall, the work presented here showcases another application of genome-engineered HSPCs as a potential one-time cure for HIV-1. Engineering CD4-expressing RBC viral traps combines promising past work utilizing CD4 as a viral decoy while building on the advantage of using cell-based carriers. More broadly, this strategy could be easily adapted to engineer expression of other viral receptors in order to treat a broad spectrum of chronic viral infections.

## Materials and Methods

### rAAV6 vector design, production, and purification

CD4 donor vectors were cloned from a gBlock Gene Fragments (Integrated DNA Technologies, IDT, San Jose, CA) using restriction enzyme digest or Gibson Assembly (New England Biolabs, Ipswich, MA) into an Adeno-associated virus, serotype 6 (AAV6) vector plasmid derived from the pAAV-MCS plasmid (Agilent Technologies, Santa Clara, CA, USA). rAAV6 vectors were either produced in-house or purchased through SignaGen Laboratories (Frederick, MD). For in-house production, HEK-293T cells were seeded in ten 15 cm^2^ dishes with 10-12×10^6^ cells per plate 24-48 hours pre-transfection. Each dish was then transfected with a standard polyethylenimine (PEI) transfection of 6 μg ITR-containing plasmid and 22 μg pDGM6 plasmid (gift from David Russell, University of Washington, Seattle, WA, USA). After 48-72 hour incubation, cells were harvested and vectors were purified using the AAVpro purification kit (cat.: 6666; Takara Bio, Kusatsu, Shiga, Japan) as per manufacturer’s instructions. Viral titers were determined using droplet digital PCR (ddPCR) to measure the number of vector genomes as previously described.^58^

### CD34^+^ HSPC isolation and culture

Human CD34^+^ HSPCs were isolated from cord blood by the Stanford Binns Program for Cord Blood Research and cultured as previously described.^59^ Samples were obtained with approval from the Stanford Institutional Review Board Committee under protocol 33813. Briefly, isolated mononuclear cells were positively selected for CD34 using the CD34+ MicroBead Kit Ultrapure (Miltenyi Biotec, San Diego, CA, cat.: 130-100-453). Cells were cultured at 1.5×10^5^–2.5×10^5^ cells/mL in CellGenix® GMP Stem Cell Growth Medium (SCGM, CellGenix, Freiburg, Germany, cat.: 20802-0500) supplemented with a human cytokine (PeproTech, Rocky Hill, NJ) cocktail: stem cell factor (100 ng/mL), thrombopoietin (100 ng/mL), Fms-like tyrosine kinase 3 ligand (100 ng/mL), interleukin 6 (100 ng/mL), streptomycin (20 mg/mL), and penicillin (20 U/mL), and 35 nM of UM171 (APExBIO, Houston, TX, cat.: A89505). Cells were cultured in a 37°C hypoxic incubator with 5% CO_2_ and 5% O_2_.

### Genome editing of HSPCs

Synthetic chemically modified sgRNAs were purchased from Synthego (Redwood City, CA) or TriLink Biotechnologies (San Diego, CA). Chemical modifications were comprised of 2′-O-methyl-3′-phosphorothioate at the three terminal nucleotides of the 5′ end and the second, third, and fourth bases from the 3’ end as described previously.^60^ The target sequence for the sgRNA targeting *HBA1* was previously published^24^ and is as follows: 5′-GGCAAGAAGCATGGCCACCG-3′. HiFi Cas9 protein was purchased from Aldevron (Fargo, ND, cat:9214). RNPs were complexed at a Cas9:sgRNA molar ratio of 1:2.5 at room temperature for 15-30 min. CD34^+^ cells were resuspended in P3 buffer (Lonza, Basel, Switzerland, cat.: V4XP-3032) with complexed RNPs and electroporated using the Lonza 4D Nucleofector and 4D-Nucleofector X Unit (program DZ-100). Electroporated cells were then plated at 2.5×10^5^ cells/mL in the previously described cytokine-supplemented media. Immediately following electroporation, AAV6 was dispensed onto cells at 1.25×10^3^-2.5×10^3^ vector genomes/cell based on titers determined by ddPCR.

### Generation of K-562 stable cell lines

K-562 cells (cat: CCL-243, ATCC) were cultured in RPMI 1640 media (cat: 16750-70, VWR) supplemented with 10% bovine growth serum (BGS), and 1% penicillin-streptomycin (RPMI complete). RNPs were complexed the same as above but using a chemically modified sgRNA targeting CCR5: 5’-TGACATCAATTATTATACAT-3’ that was previously published.^34^ 1×10^6^ K-562 cells were resuspended in P3 buffer (Lonza, Basel, Switzerland, cat.: V4XP-3032) with complexed RNPs and 250 fmol of donor construct plasmid. Cells were then electroporated using the Lonza 4D Nucleofector and 4D-Nucleofector X Unit (program FF-120) and then replated in 2 mL RPMI complete media. Following recovery and expansion, cells were sorted for GFP using a FACSAria II SORP (BD), twice for purity. Cells were then stained with a CD4-PE (clone: RPA-T4, BioLegend, San Diego, CA, cat: 300508) antibody and analyzed via flow cytometry for confirmation of CD4 expression.

### Allelic modification analysis by ddPCR

Genomic DNA was extracted from cells using QuickExtract DNA extraction solution (Biosearch Technologies, Hoddesdon, UK, cat.: QE09050). To quantify knock-in alleles via ddPCR, we employed *HBA1* specific in-out PCR primers and a probe corresponding to the expected knock-in event (1:3.6 primer to probe ratio). We also used an established genomic DNA reference at the *CCRL2* locus.^61^ The ddPCR reaction was prepared and underwent droplet generation following the manufacturer’s instructions with a Bio-Rad QX200 ddPCR machine (Bio-Rad, Hercules, CA). Thermocycler settings were as follows: 95°C (10 min, 1°C/s ramp), 94°C (30 s, 1°C/s ramp), 57.7°C (30 s, 1°C/s ramp), 72°C (2 min, 1°C/s ramp), return to step 2 for 50 cycles, and 98°C (10 min, 1°C/s ramp). Analysis of droplet samples was then performed using the QX200 Droplet Digital PCR System (Bio-Rad). We divided the copies/μL to determine the frequency of HDR (%): HDR (FAM) / REF (HEX). The following primers and probes were used in the ddPCR reaction:

*CD4 constructs:*

Forward Primer (FP): 5’-TGCTGTCCGAGAAGAAAACC-3’

Reverse Primer (RP): 5’-TAGTGGGAACGATGGGGGAT-3’

Probe: 5’-6-FAM/TGCTGGAGTGGGACTTCTCT/3IABkFQ-3’

*GPA constructs:*

Forward Primer (FP): 5’-GAAATCGAGAACCCCGAGAC-3’

Reverse Primer (RP): 5’-TAGTGGGAACGATGGGGGAT-3’

Probe: 5’-6-FAM/ TGCTGGAGTGGGACTTCTCT/3IABkFQ-3’

*tEPOR constructs:*

FP: 5’-TCTGCTGCCAGCTTTGAGTA-3’

RP: 5’-GCTGGAGTGGGACTTCTCTG-3’

Probe: 5’-6-FAM/ACTATCCTGGACCCCAGCTC/3IABkFQ-3’

*CCRL2 (reference):*

FP: 5’-GCTGTATGAATCCAGGTCC-3’,

RP: 5’-CCTCCTGGCTGAGAAAAAG −3’

Probe: 5’-HEX/TGTTTCCTC/ZEN/CAGGATAAGGCAGCTGT/3IABkFQ −3’

### Differentiation of HSPCs into erythrocytes

Two to three days following editing of CD34^+^ HSPCs, a 14-day in vitro differentiation was performed in supplemented SFEMII medium as previously described.^24,33,62^ For the first phase of differentiation, SFEMII base medium was supplemented with 100 U/mL penicillin–streptomycin, 10 ng/mL SCF (PeproTech, Rocky Hill, NJ), 1 ng/mL IL-3 (PeproTech, Rocky Hill, NJ, USA), 3 U/mL EPO (eBiosciences, San Diego, CA), 200 μg/mL transferrin (Sigma-Aldrich, St. Louis, MO), 3% human serum (heat-inactivated from Sigma-Aldrich, St. Louis, MO or Thermo Fisher Scientific, Waltham, MA), 2% human plasma (isolated from umbilical cord blood provided by Stanford Binns Cord Blood Program), 10 μg/mL insulin (Sigma-Aldrich, St. Louis, MO), and 3 U/mL heparin (Sigma-Aldrich, St. Louis, MO). Cells were cultured in the first phase of medium for seven days at 1×10^5^ cells/mL. In the second phase of medium, days 7-10, cells were maintained at 1×10^5^ cells/mL, and IL-3 was removed from the culture. In the third phase of medium, days 11-14, cells were cultured at 1×10^6^ cells/mL, with a transferrin increase to 1 mg/mL. When cells were kept past day 14, the third phase of medium was used and cell concentration was adjusted to 5×10^6^ cells/mL.

### Colony forming unit (CFU) assay

2-3 days post-electroporation 500 HSPCs were plated in each well of a SmartDish 6-well plate (STEMCELL Technologies, Vancouver, Canada, cat.: 27370) containing MethoCult H4434 Classic (STEMCELL Technologies, cat.: 04444). After 12-14 days, the wells were imaged using the STEMvision Hematopoietic Colony Counter (STEMCELL Technologies). Colonies were counted and scored with manual correction to determine the number of BFU-E, CFU-GM, and CFU-GEMM colonies.

### Immunophenotyping of differentiated red blood cells

Differentiated erythrocytes were analyzed using flow cytometry using a previously published panel of RBC markers.^24^ Cells were stained for 15-30 min at 4°C with the following antibodies: hCD45-V450 (clone HI30, cat: 560367 BD Biosciences, San Jose, CA, USA), CD34-APC (clone 561, cat: 343510 BioLegend, San Diego, CA), CD71-PE-Cy7 (clone OKT9, cat: 25-0719-42, Invitrogen, Santa Clara, CA, USA), CD235a-PE (GPA) (clone GA-R2, cat: 12-9987-82, BD Biosciences), CD4-FITC (clone RPA-T4, cat: 300506, BioLegend), and Ghost Dye Red 780 viability dye (Tonbo Biosciences, San Diego, CA). To analyze more mature RBC markers, cells were first blocked for non-specific binding with human FcR Blocking Reagent (Miltenyi Biotec, cat.: 130-059-901) for 10 min at room temperature and then stained for 20-30 min at 4°C with the following antibody cocktail: CD71-PE-Cy7 (clone OKT9, cat: 25-0719-42, Invitrogen, Santa Clara, CA, USA), and CD235a-PE (GPA) (clone GA-R2, cat: 12-9987-82, BD Biosciences), CD49d-BV711 (α4-integrin) (clone 9F10, cat: 304331, BioLegend, San Diego, CA), CD233-APC (Band3) (clone BRIC 6, cat: 9439, Internation Blood Group Reference Laboratory, Filton, Bristol, UK), CD4-FITC (clone RPA-T4, cat: 300506, BioLegend), and Ghost Dye Red 780 viability dye (Tonbo Biosciences, San Diego, CA, USA). Samples were analyzed on a FACSAria II SORP (BD). Data was subsequently analyzed using FlowJo (v.10.10.0).

### CD4^+^ T cell isolation and culture

Peripheral blood mononuclear cells (PBMCs) were isolated from LRS chambers obtained from healthy donors using a Ficoll density gradient. Subsequently, CD4^+^ cells were isolated by negative selection using CD4^+^ T Cell Isolation Kit, human (Miltenyi, cat: 130-096-533). T cells were cultured at 1×10^6^ cells/mL in X-VIVO15 with gentamicin (Lonza), 10% bovine growth serum (BGS), and 100 IU mL^−1^ human IL-2 (Peprotech) at 37 °C with 5% CO2 and ambient O2. For the stimulated condition, cells were activated for 3 days with anti-CD3/anti-CD28 paramagnetic beads (Dynabeads Human T Cell Activator, Gibco, cat: 11161D).

### Modified TZM-bl neutralization assay

The modified TZM-bl HIV-1 pseudotype neutralization assay was adapted from a previously described protocol.^11^ TZM-bl cells were obtained through the NIH AIDS Reagent Program (cat.: 8129) and cultured in DMEM with 10% BGS, and 1% penicillin-streptomycin (complete DMEM) at 37°C, 5% CO2, and ambient oxygen levels. HIV-1 env pseudotyped lentivirus was produced as previously described using a pNL4-3ΔEnvGFP backbone plasmid and *env* plasmids provided by the NIH AIDS Reagents Program (cat: 11100 and cat: 12670, respectively).^35,63^ Differentiated erythrocytes were resuspended in complete DMEM with 10 µg/mL polybrene (cat: TR1003G, Fisher Scientific), serial dilutions were performed, and dilutions were moved to a 48 well plate. HIV-1 pseudovirus was then added to each well and plate was incubated for 4 hours while shaking at 600 rpm at room temperature. 1×10^4^ TZM-bl cells were plated in black-walled, clear-bottom 96-well plates (Corning, Corning, NY, cat.: 07-200-565). RBCs were then spun down at 500 g for 10 min in a 96-well U-bottom plate (in duplicate) and supernatant was then moved to the well of the black-walled, clear-bottom 96-well plates containing TZM-bl cells. Cells were incubated for 48 hours then measured for luciferase signal corresponding to infection. Briefly, cells were lysed with Reporter Lysis Buffer (Promega, Madison, WI, cat.: E3971) and freeze-thawed at −80°C. Lysed samples were read for relative luminescence units (RLU) using a Synergy H1 plate reader (BioTek, Winooski, VT) that injected luciferin solution consisting of the following: 200 mM Tris [pH 8], 10 mM MgCl2, 300 μM ATP, Firefly Luciferase Signal Enhancer (Thermo Fischer Scientific, cat. 16180), and 150 μg/mL d-luciferin (Biosynth Chemistry & Biology, Staad, Switzerland, cat.: L8220). Percent infection was determined by normalizing RLU values to the RLU of Mock-edited RBC wells using GraphPad Prism v10.

### Statistical analysis

GraphPad Prism v10 software was used for all statistical analysis.

## Data availability statement

The main data supporting the results in this study are available within the article and its supplementary figures. Sequences of the sgRNA and ddPCR primers and probes are provided in Materials and Methods section. Source data for the figures are provided with this paper. All data generated in this study are available from the corresponding authors upon reasonable request.

## Acknowledgements

We thank the NIH AIDS Reagent Repository for providing the *env* plasmids for the HIV-1 pseudoviruses. We thank the Stanford Institute for Stem Cell Biology and Regenerative Medicine FACS Core for their support and access to the flow cytometry machines. We thank the Binns Program for Cord Blood Research for providing purified cord blood HSPCs. S.E.L was supported by the Stanford Medical Scholars Research Program, the American Society of Hematology Minority Medical Student Award Program, the Stanford Medical Scientist Training Program, and NIH F30 Individual Predoctoral Fellowship (1F30HL178496-01). W.N.F was supported by the Stanford T32 Graduate Training Program in Stem Cell Biology and Regenerative Medicine (5T32GM119995) and the Blavatnik Family Foundation as a Blavatnik Fellow. F.K.E. was supported by the Stanford Knight-Hennessy Scholarship, Paul And Daisy Soros Fellowship for New Americans, and the Hertz Fellowship. A.M.D. was supported by NIH F32 Individual Postdoctoral Fellowship (1F32HL154667-01)) and NIH K99/R00 Pathway to Independence Award (1K99HL172253-01). M.H.P. was supported by the Sutardja Chuk Professorship in Definitive and Curative Medicine and the Laurie Kraus Lacob Translational Medicine endowment.

## Author Contributions

S.E.L, W.N.F., K.B.E, N.A.A., N.M.J., H.Y.G., F.K.E., and A.M.D. contributed to experimental design, performance, and analysis. S.E.L, W.N.F., K.B.E., and M.H.P. wrote the draft and finalized versions of the manuscript with input from other authors. M.H.P. supervised the project.

---

**Supplementary Figure 1:**
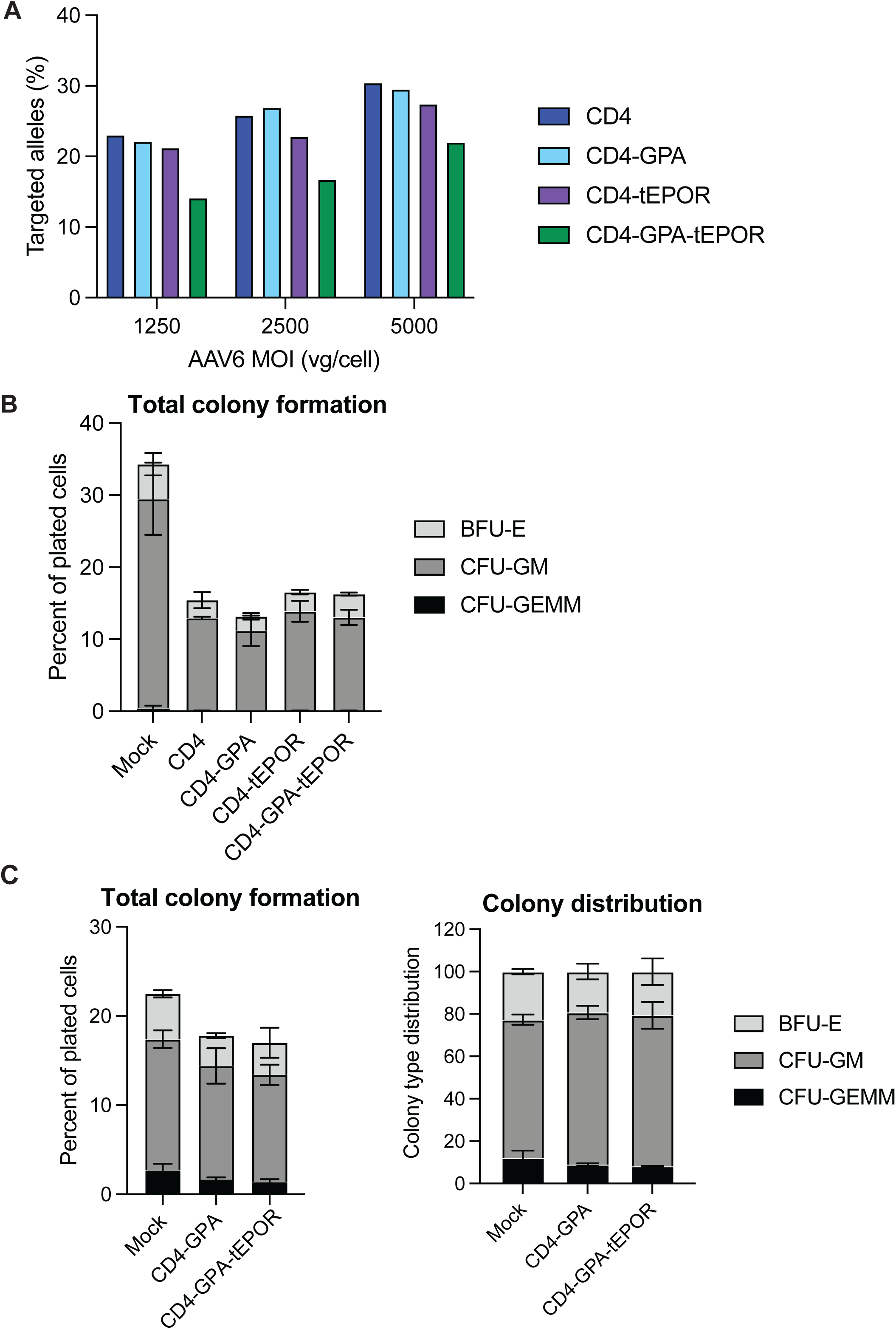
Further characterization of editing of HSPCs with CD4-expression cassettes. **A.** Targeted integration of CD4-expression cassettes in CD34^+^ HSPCs at three different amounts of AAV6 per cell, 1250, 2500 and 5000 vg/cell. n=1. **B.** Total colony formation as a percentage of plated cells (500 cells plated per condition). Bars represent mean +/− SD. n=3 biological donors.. **C.** Total colony formation as a percentage of plated cells (500 cells plated per condition) (left) and colony distribution (right) of an edited HSPC donor edited with higher-quality, purified by cesium chloride (CsCl) gradient ultracentrifugation, AAV6. Bars represent relative frequency of each colony type: CFU-GEMM, CFU-GM, BFU-E. n=1 donor. All bars represent mean +/− SD.

**Supplementary Figure 2:**
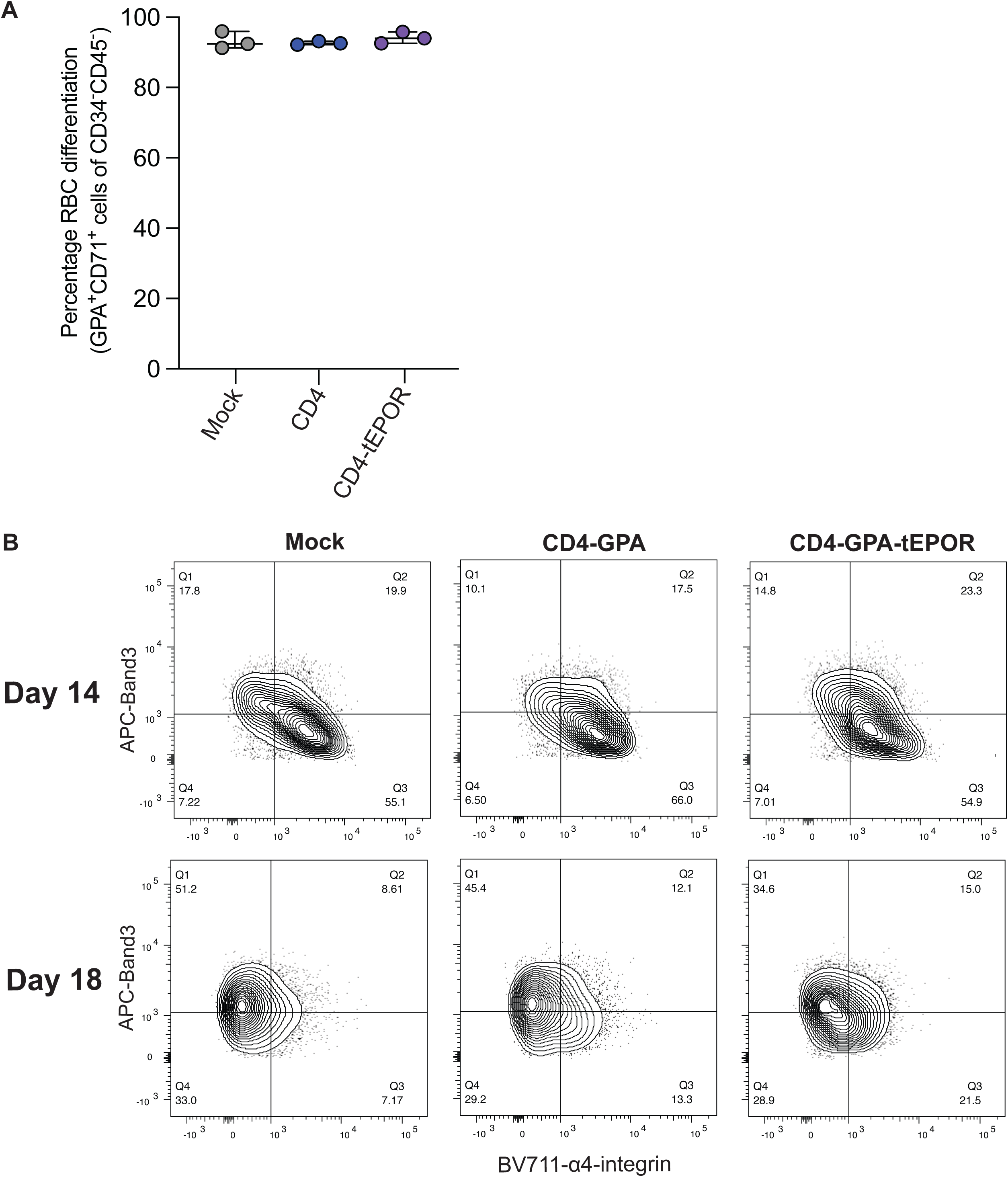
Immunophenotyping of differentiated CD4-expressing RBCs. **A.** Percentage GPA^+^CD71^+^ cells of CD34^-^CD45^-^ cells on day 14 of RBC differentiation. Lines represent median +/− 95% confidence interval. n=3 independent biological donors. **B.** Flow cytometry plots of expression of Band3 in differentiated Mock, CD4-GPA, and CD4-GPA-tEPOR RBCs on day 14 and 18 of RBC differentiation. Plots shown as percent of live single CD71^+^ cells. n=1.

**Supplementary Figure 3:**
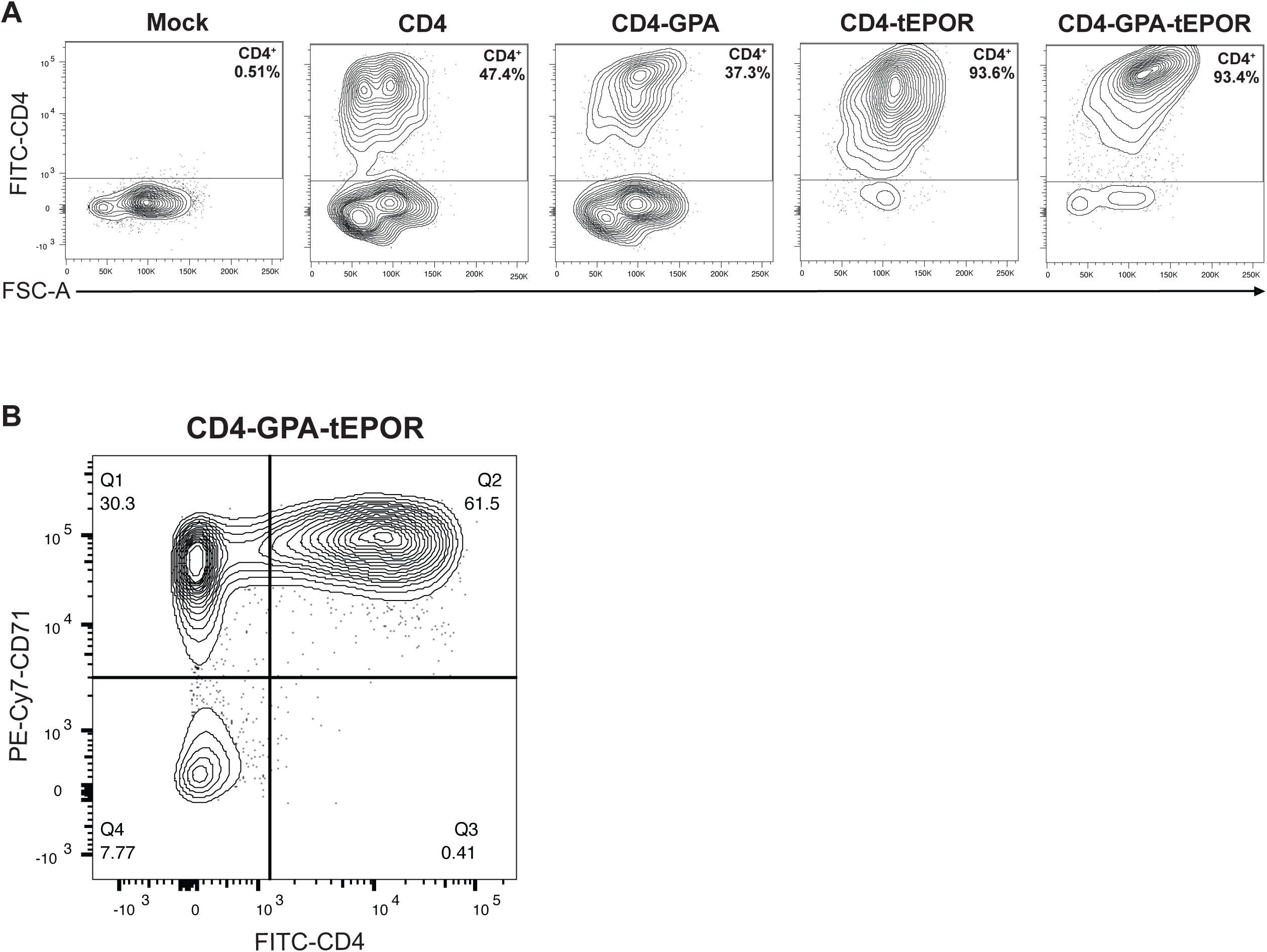
Flow cytometry plots of CD4 expression in differentiated RBCs. **A.** Representative flow cytometry plots of CD4 expression on day 14 of RBC differentiation. **B.** Representative flow cytometry plot of CD4-GPA-tEPOR expressing cells on day 7 of differentiation showing CD4 expression restricted to cells expressing RBC marker CD71. All plots gated on live single cells.

**Supplementary Figure 4:**
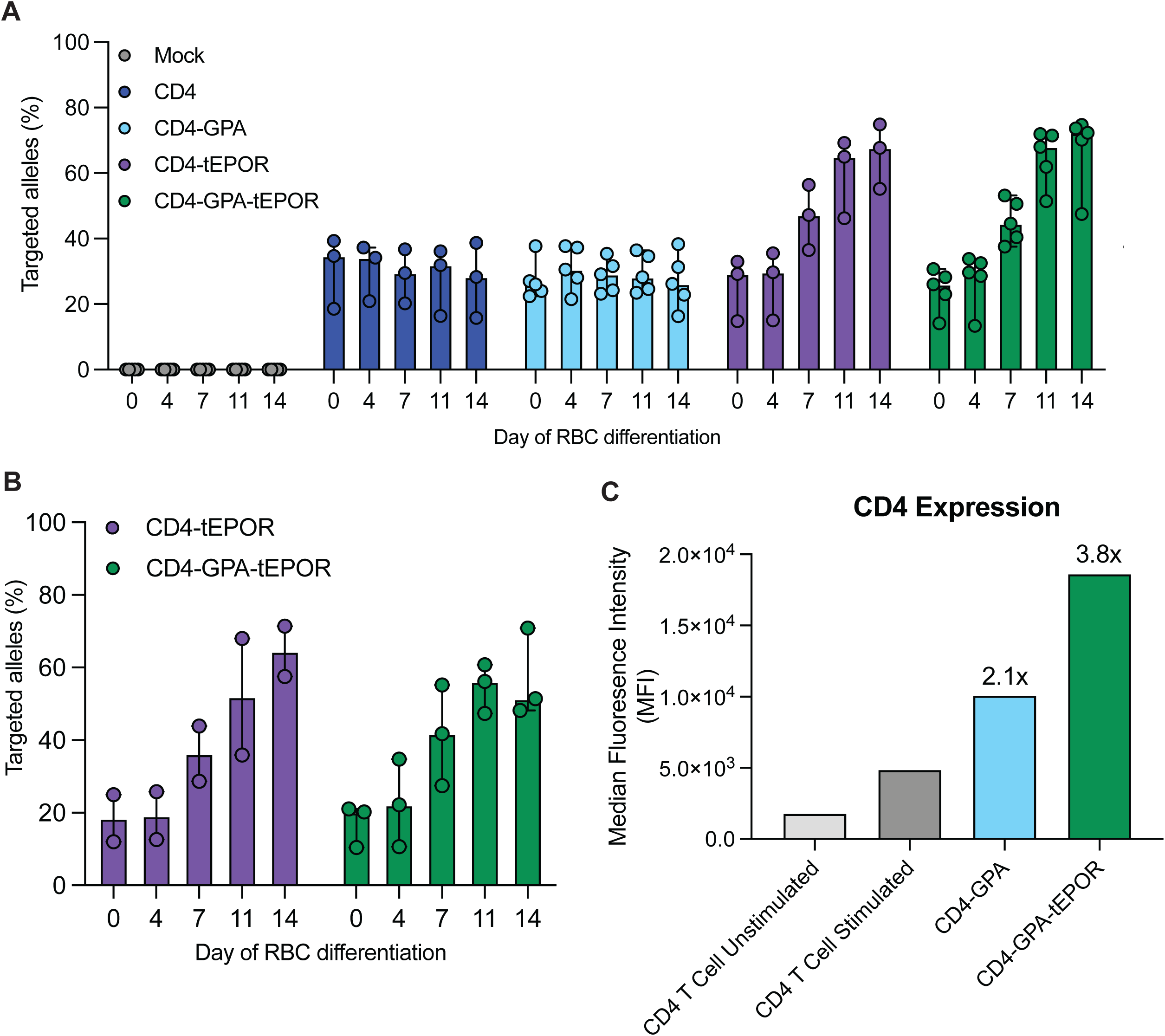
Targeted integration of edited HSPCs throughout differentiation and relative CD4 expression. **A.** Percentage of targeted integration of CD4 cassettes over the course of RBC differentiation. n=5 independent biological donors for Mock, CD4-GPA, CD4-GPA-tEPOR, n=3 for CD4 and CD4-tEPOR. Bars represent median +/− 95% confidence interval. **B.** Percentage of targeted integration of CD4-tEPOR and CD4-GPA-tEPOR during RBC differentiation of HSPCs targeted with lower amount of AAV6 (1250 vg/cell). n=2 independent biological donors. Bars represent median +/− 95% confidence interval. **C.** Relative CD4 expression from day 14 differentiated CD4-GPA and CD4-GPA-tEPOR RBCs compared with stimulated and unstimulated CD4^+^ T cells on day 3 post-thaw with or without CD3/CD28 stimulation. Cells gated on live single cells. Fold increase in MFI compared to stimulated CD4 T cells shown above bars for CD4-GPA and CD4-GPA-tEPOR conditions. n=1 biological donor.

**Supplementary Figure 5:**
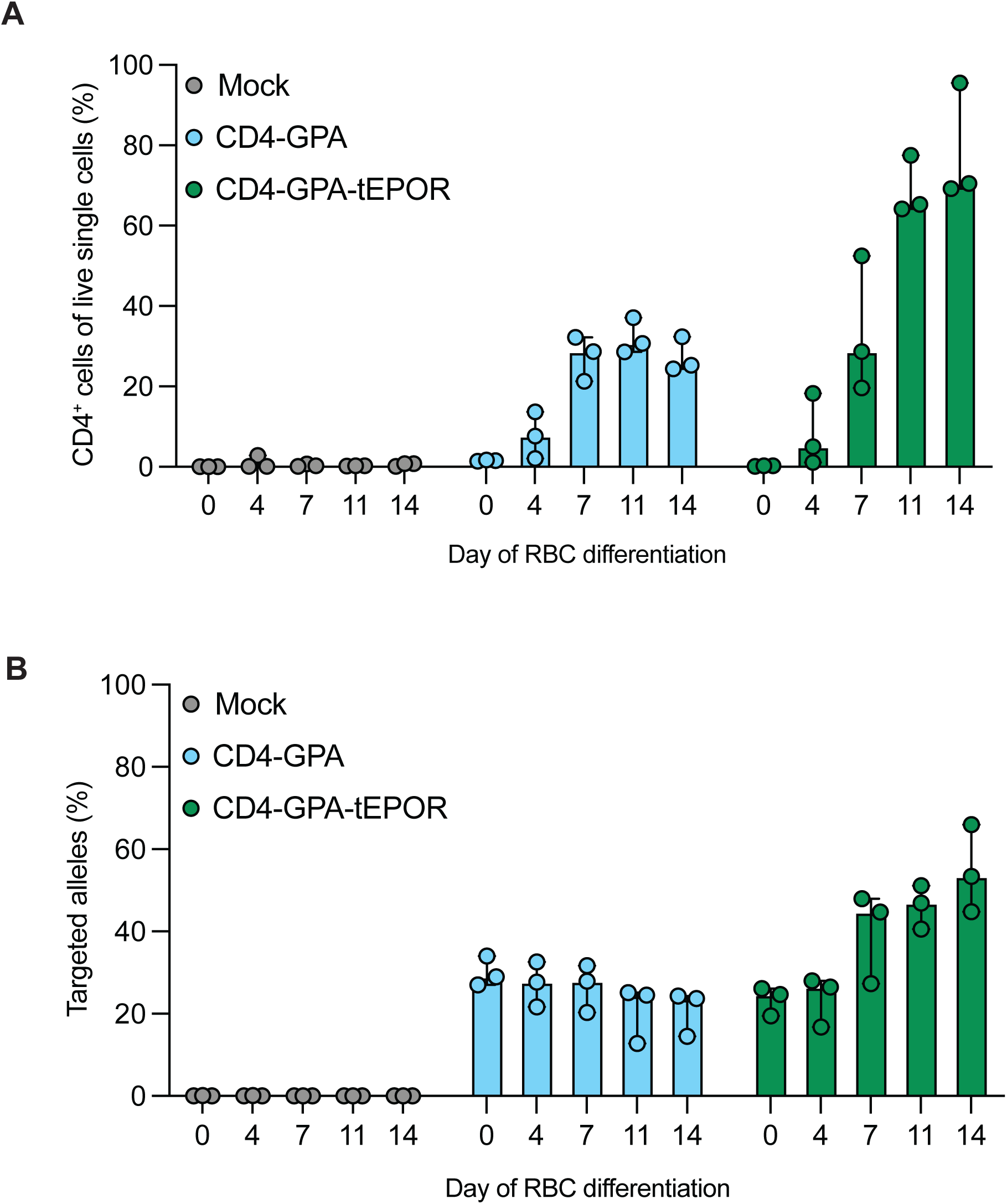
CD4 expression and targeted integration during RBC differentiation of CD4 expressing RBCs used in TZM-bl neutralization assay. **A.** Percentage of CD4^+^ cells of live single cells on each day of RBC differentiation for donors used in TZM-bl assay (Figure 4). n=3 independent biological donors. **B.** Targeted integration of CD4 expression cassettes during RBC differentiation for donors used in TZM-bl assay. n=3. All bars represent median +/− 95% confidence interval.

